# Infinite re-reading of single proteins at single-amino-acid resolution using nanopore sequencing

**DOI:** 10.1101/2021.07.13.452225

**Authors:** Henry Brinkerhoff, Albert S. W. Kang, Jingqian Liu, Aleksei Aksimentiev, Cees Dekker

## Abstract

As identifying proteins is of paramount importance for cell biology and applications, it is of interest to develop a protein sequencer with the ultimate sensitivity of decoding individual proteins. Here, we demonstrate a nanopore-based single-molecule sequencing approach capable of reliably detecting single amino-acid substitutions within individual peptides. A peptide is linked to a DNA molecule that is pulled through the biological nanopore MspA by a DNA helicase in single amino-acid steps. The peptide sequence yields clear stepping ion current signals which allows to discriminate single-amino-acid substitutions in single reads. Molecular dynamics simulations show these signals to result from size exclusion and pore binding. Notably, we demonstrate the capability to ‘rewind’ peptide reads, obtaining indefinitely many independent reads of the same individual molecule, yielding virtually 100% read accuracy in variant identification, with an error rate less than 10^−6^. These proof-of-concept experiments constitute a promising basis for developing a single-molecule protein sequencer.

**One-sentence summary:** This paper presents proof-of-concept experiments and simulations of a nanopore-based approach to sequencing individual proteins.

---

Over the past half century, advances in DNA sequencing technology have revolutionized biology. While the genome is a key source of basic information, it has become clear that splicing, transcriptional variants, and post-translational modifications lead to an enormous diversity of proteins, and neither the DNA genotype nor the RNA transcriptome can fully describe the protein phenotype(*1*). With nucleic acid sequencing now a solved problem, the next wave of research in fundamental biology, biotechnology, and bioinformatics will revolve around sequencing the enormous and dynamic variations in the proteome.

Sequencing proteins is a costly and time-consuming process that faces significant intrinsic limitations(*2, 3*). Protein sequencing and detection of post-translational modifications (PTMs) by mass spectrometry, the current gold standard, requires extensive sample preparation and large sample sizes, and possesses a limited dynamic range with respect to protein concentrations, limitations that severely restrict its application to biological and clinical problems. A robust method for sequencing proteins and detecting PTMs at the single-molecule level would be revolutionary for proteomics research(*4*), allowing biologists to quantify low-abundance proteins as well as distributions and correlations of PTM patterns, all at a single-molecule and single-cell level. In this work, we provide proof-of-concept data for a nanopore-sequencing-based approach that is able to discriminate single peptides at single-amino-acid sensitivity with unprecedented fidelity and potential for high throughput.

Recently, biological nanopores have been used as the basis of a single-molecule DNA sequencing technology(*5*) that is capable of long reads(*6, 7*) and detection of epigenetic markers(*8–10*) in a portable platform with minimal cost. In such experiments, single-stranded DNA is slowly moved step-by-step through a membrane channel embedded in a thin membrane, partially blocking an electrical current carried by ions through the nanopore. The DNA stepping is accomplished using a DNA-translocating motor enzyme, such as a polymerase or a helicase, which slowly walks along the DNA, moving the DNA through the pore in discrete steps and yielding a series of steps in the ion current. Each ion current level characterizes the bases residing in the pore at that step, and the sequence of levels can be decoded into the DNA base sequence.

It has been hypothesized that nanopores can also be used for protein fingerprinting or sequencing(*11–15*). Methods in which small peptide fragments freely translocate through a pore have shown sensitivity to single amino acids(*16–18*), but lack a method for determining the order of amino acids and reconstructing the sequences of single proteins. Enzyme-controlled methods using a protease to pull a peptide through a nanopore have yielded signals that effectively distinguished between different peptides(*19*), but were difficult to interpret, in part due to the irregular stepping behavior of protein unfoldases(*20*) compared to the very regular single(*21, 22*)- or half(*23*)-nucleotide steps taken by DNA-translocating motors.

Here, we apply the exquisite stepwise control of a DNA-translocating motor to pull a peptide through a nanopore. As illustrated in Fig. 1, we developed a system in which a DNA-peptide conjugate is pulled through a biological nanopore by a helicase that is walking on the DNA section. The conjugate strand consisted of an 80-nt DNA strand that is covalently linked to a 26-amino-acid synthetic peptide by means of a DBCO click linker on the 5’ end of the DNA connecting to an azide modification at the C-terminus of the peptide (Supplementary Fig. S1). A negatively charged peptide sequence (mostly aspartic acid and glutamic acid residues) was chosen such that the electrophoretic force assisted in pulling the peptide into the pore. The main DNA construct (Fig. 1A) consisted, first, of a template strand with a 3’ overhang acting as a helicase loading site. A complementary DNA strand blocks the procession of the helicase until the construct is pulled into the pore(*22, 23*), which shears off the complement and allows the helicase to begin stepping the analyte back up through the pore (Fig. 1B). The complementary strand has a poly-T_20_ tail with a 3’ cholesterol, which associates with the bilayer and increases the concentration of analyte near the pore, improving the event rate(*7, 24, 25*). For sequencing, we used the mutant nanopore M2 MspA(*26*) with a cup-like shape(*27*) that separates the anchoring enzyme by ~10 nm from the constriction of the pore where the blockage of ion current occurs (*28*). For the DNA-translocating motor enzyme, we used Hel308 DNA helicase because its observable half-nucleotide ~0.33 nm steps, when pulling ssDNA through MspA(*23*), are approximately the same as single-amino acid steps.

**Figure 1:**
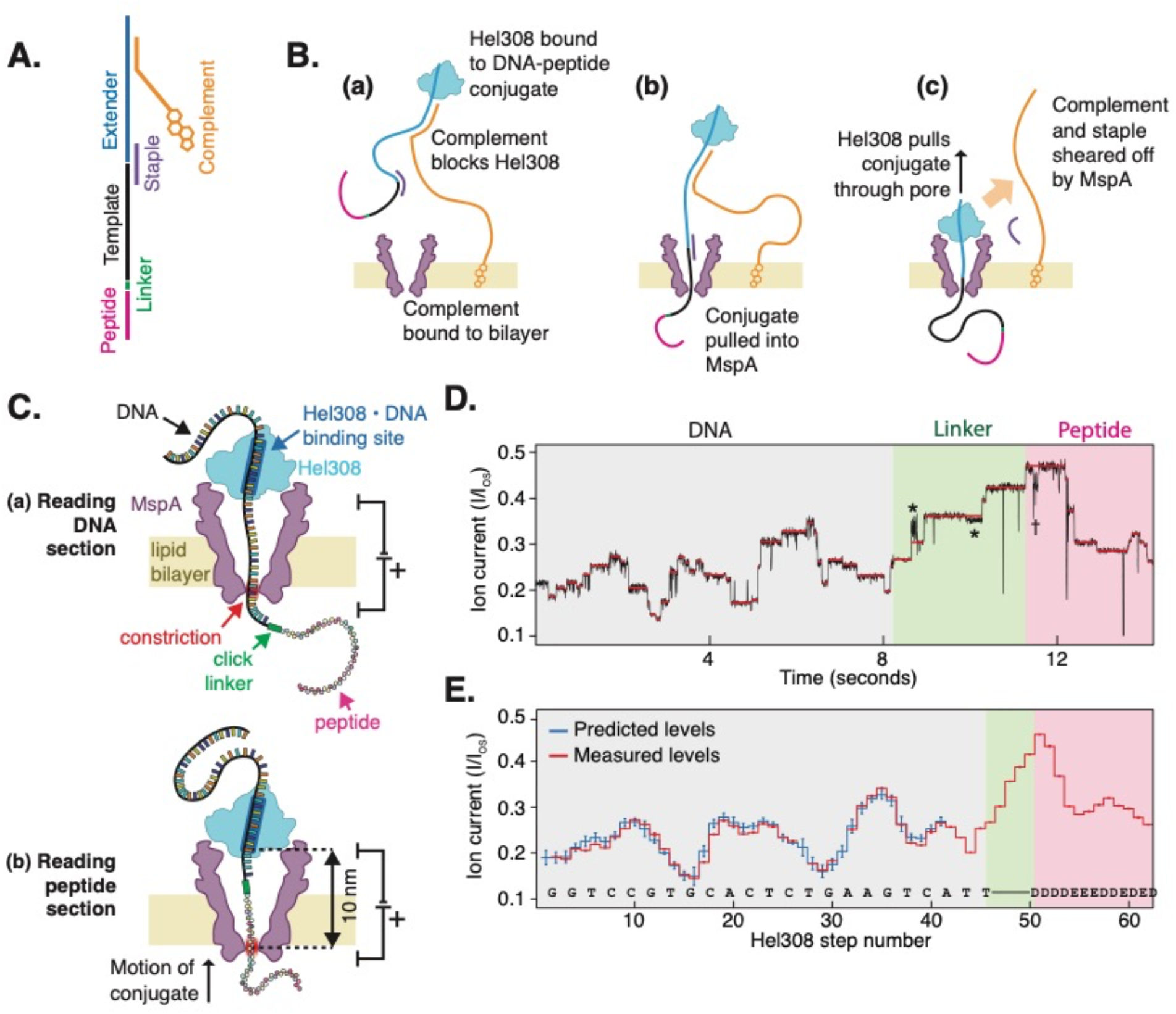
Reading peptides with a nanopore sequencer. (A) The DNA-peptide conjugate construct consists of a peptide (pink) attached via a click linker (green) to an ssDNA strand (black). This DNA-peptide conjugate is extended with a typical nanopore sequencing adaptor comprised of an extender that acts as a site for helicase loading (blue) and a complementary oligo with a 3’ cholesterol modification (gold). (B) The cholesterol associates with the bilayer as shown in (a), increasing the concentration of analyte near the pore. The complementary oligo blocks the helicase, until it is pulled into the pore (b), causing the complementary strand to be sheared off (c), whereupon the helicase starts to step along DNA. (C) As the helicase walks along the DNA, it pulls it up through the pore, resulting in (a) a read of the DNA portion followed by (b) a read of the attached peptide. (D) Typical nanopore read of a DNA-peptide conjugate (black), displaying clear step-like ion currents (identified in red). The asterisks * indicate a spurious level not observed in most reads and therefore omitted from further analysis. The dagger † indicates a helicase backstep. (E) Consensus sequence of ion current steps (red), which for the DNA section is closely matched by the predicted DNA sequence (blue). The linker and peptide sections are identified by counting half-nucleotide steps over the known structural length of the linker. Error bars in the measured ion current levels are errors in the mean value, often too small to see. Error bars in the prediction are standard deviations of the ion current levels that were used to build the predictive map in previous work(*29*).

As shown in Fig, 1C, we find that, similarly to nanopore-based DNA sequencing, ratcheting a peptide through the nanopore generates a distinct step-like pattern in the ion current. Durations of ion current steps vary from read to read, but the sequence of levels, which encodes sequence information, is highly reproducible (Supplementary Fig. S3). Each step of the ion current corresponds to a single half-nucleotide step of the helicase. The progression of current level steps was accurately identified using custom software (Supplementary Note 5) and further analysis was performed on the sequence of the median values of ion current for each step (Fig. 1D).

This sequence of ion current levels first closely tracks the sequence expected for the template strand of DNA, which can be predicted using a DNA-sequence-to-ion-current map developed previously(*7, 29*). After the end of the DNA has crossed to the *cis* side of MspA’s constriction, we continue to observe stepping over the linker (a length of ~2 nm, or 6 Hel308 steps), and subsequently over the peptide. The stepping of the peptide through the MspA constriction produced well-distinguishable ion current steps, much like those from DNA, but with a higher average ion current. The higher current is expected because the smaller amino acids are expected to block the ion current less than DNA nucleotides. While individual reads may contain a varying number of steps due to helicase backstepping and errors in step segmentation, we identified these features by cross-comparison of several independent reads, producing a “consensus” ion current sequence free of helicase mis-steps or step-segmentation errors. By counting the steps in these consensus sequence traces, we determined the parts of the traces that correspond to the linker (the first 6 steps after the DNA) and the peptide (all steps thereafter) in the MspA constriction. We confirmed this analysis by altering the peptide sequence at a selected site and observing the location of the resulting change in the ion current stepping sequence, as discussed below.

Remarkably, our approach allows us to discriminate peptide variants that differ by only a single amino acid. We obtained reads (N=211) of three different DNA-peptides in nineteen different pores, where the peptide sequences consisted of a mixture of negatively charged aspartic acid (D) and glutamic acid (E) residues, with a single variation, i.e., aspartic acid (D), glycine (G), or tryptophan (W), placed four amino acids away from the C-terminus that connects to the linker, see Supplementary Table S1 for full sequences. The three variants showed a reproducible difference at the site of the substituted amino acid, which can be seen by comparing the consensus sequences of ion current levels, see Figs. 2A and B. As is typical of nanopore experiments, a single-site variation was found to affect several ion current steps, because multiple amino acids around the pore constriction of MspA affect the ion current blockage level (*7, 21*) due to the finite constriction height and stochastic displacements of the strand up and down through the nanopore(*30*). The center of the differing region in the ion current sequence is at the expected site: about 10 helicase steps away from the end of the DNA section (6 half-nucleotide steps for the linker and 4 more along the peptide to the variant site). The signals vary by several standard deviations over multiple sequential levels, clearly demonstrating that variations as small as a single-amino-acid substitution can be resolved. The differences of the ion currents for the W- and G- substituted variants from the D-substituted variant (Fig. 2B) show an interesting behavior: when the G amino acid, which has merely a hydrogen atom as a side chain, occupies the nanopore constriction, we see higher ion current levels, as expected from a smaller amino acid volume. But when the bulky W variant moves through the constriction, the ion current, counterintuitively, first decreases and then increases relative to the medium-sized D variant.

**Figure 2:**
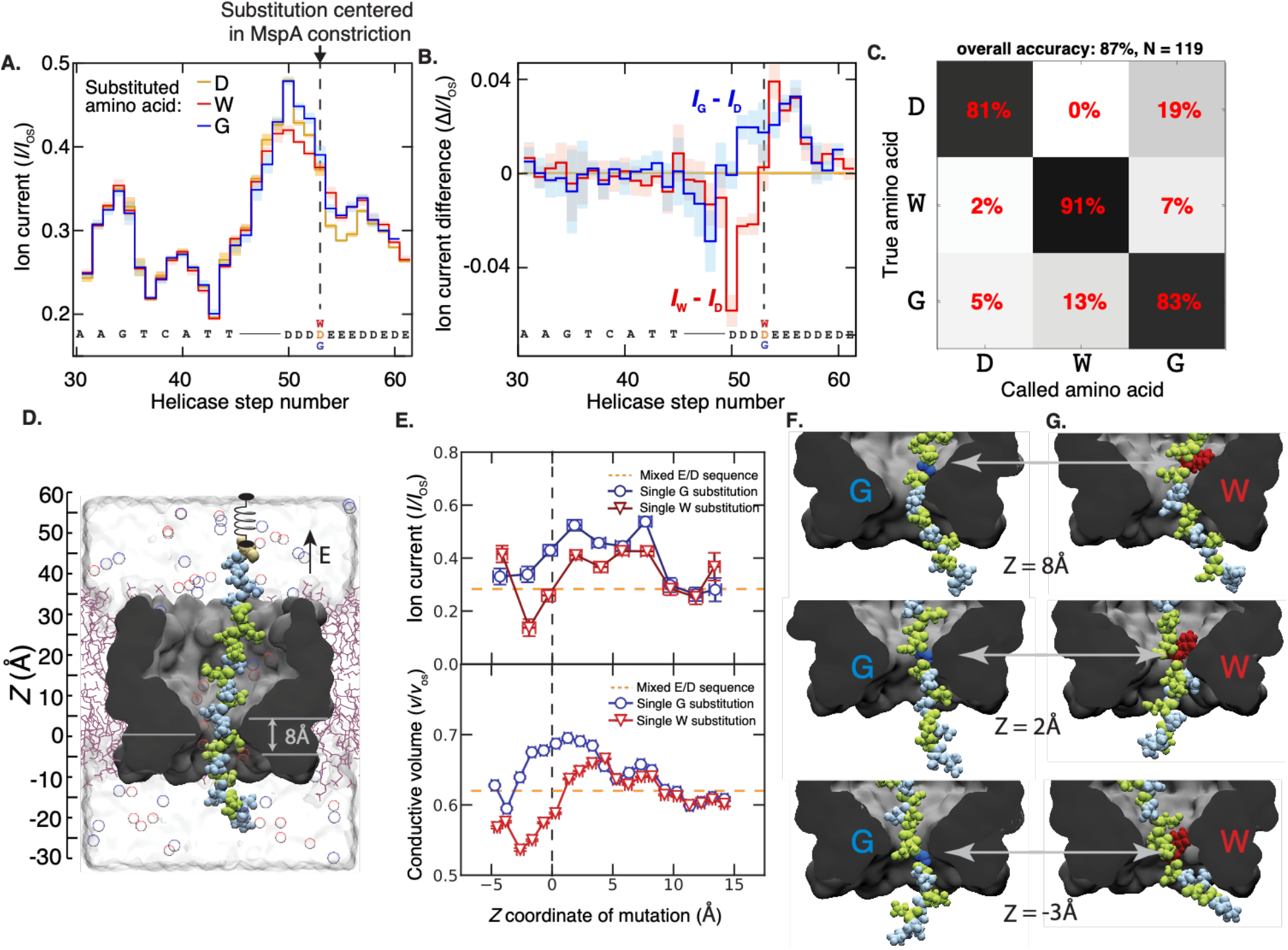
Detection of single amino acid substitutions in single peptides. (A) Consensus ion current sequences for each of the three measured variants (D, gold; W, red; G, blue), which differ significantly at the site of the amino acid substitution. (B) Difference in ion current between the W (red) and G (blue) variants and the D variant. Error bars are standard deviations. (C) Confusion matrix showing error modes of a blind classifier in identifying variants of reads, demonstrating an 87% sequencing accuracy. (D) All-atom model where a reduced-length MspA pore (grey) confines a polypeptide chain (Glu: green, Asp: light blue; Cys: beige). The top end of the peptide is anchored using a harmonic spring potential, representing the action of the helicase at the rim of a full-length MspA. Water and ions are shown as semitransparent surface and spheres, respectively. (E) Top: Ionic current in MspA constriction versus *z* coordinate of the mutated residue backbone from MD simulations. Bottom: Fraction of nanopore construction volume available for ion transport. Vertical and horizontal error bars denote standard errors and standard deviations, respectively. (G,H) Representative molecular configurations observed in MD simulations of peptide variants. Glycine and tryptophane residues are shown in dark blue and red, respectively. Significant peptide/pore surface interactions are observed.

To understand the origin of these patterns, we performed all-atom molecular dynamics simulations measuring the ion current with peptide variants at varying positions within the MspA constriction. In a typical simulation setup, a polypeptide chain was threaded through a reduced-length model of MspA nanopore that was embedded in a lipid bilayer and surrounded by 0.4M KCl electrolyte (Fig. 2C). Peptides with either one W or G substitution in a mixed D/E sequence were examined under a +200 mV bias at various locations relative to the MspA constriction (see Supplementary Note 11 and Supplementary Figs. S8-S11 for details) Two patterns of ionic current blockades resulted, Fig. 2E (top panel), that, gratifyingly, matched the counterintuitive blockade current patterns that were experimentally measured for G and W substitutions (cf. Fig. 2A,B). Furthermore, the ion current correlates with the nanopore constriction volume that is available for ion transport near the pore mouth, Fig. 2E (bottom panel), with the latter quantity being more accurately characterized by the all-atom MD method (*30*). In the case of a glycine G residue, its upward motion is seen to be accompanied by an increase of the nanopore volume (Fig. 2E, bottom), that subsides as the residue leaves the nanopore constriction (Fig. 2F), in sync with the blockade current (Fig. 2E, top). A tryptophan W residue, however, reduces the nanopore constriction volume when it is located below the constriction (Fig. 2E, top), but increases the volume at and above the constriction. The latter counterintuitive effect can be traced back to a binding of the W side chain to the nanopore surface above the constriction (Fig. 2G). Thus, a glycine substitution merely increases the nanopore volume as the residue passes through the constriction, whereas the tryptophan residue can either decrease the volume when its side chain enters the constriction or increase the volume, when its side chain enters the constriction or subsequently increase the volume when its side chain binds to the inner nanopore surface.

To quantitatively assess the distinguishability of peptide variants, we computed a so-called confusion matrix, Fig. 2D. Using a hidden Markov model (Supplementary Note 7), we quantified the relative likelihoods of the alignments to the three consensus sequences for 119 reads withheld from the consensus sequence generation, finding that we can identify the correct variant with, on average, 87% accuracy. This high level of accuracy compares favorably to early nanopore DNA sequencing experiments, which identified single-nucleotide variants with significantly lower accuracy(*7*). Still, the limited single-read accuracy is an ongoing challenge in nanopore sequencing approaches, requiring the implementation of strategies to increase sequencing fidelity to acceptable levels (*29, 31, 32*). The largest error modes in DNA nanopore sequencing are due to random effects as enzymes step stochastically both forwards and backwards, and sometimes step too quickly to be clearly resolved, resulting in incorrect step identifications. This introduces a significant amount of random error to single reads, which in DNA sequencers is typically addressed by obtaining 20x coverage or more, i.e., where 20 or more independent reads are averaged. However, for a truly single-molecule technology, single-read accuracy is essential.

Strikingly, however, our nanopore protein sequencing approach allows us to increase identification fidelity dramatically to 100% by obtaining indefinitely many independent re-readings of the same individual molecule with a succession of controlling helicases, eliminating the random errors that lead to inaccuracies in nanopore sequencing. At a very high concentration of helicase, on the order of 1 μM, the DNA in the pore will have a second helicase queued up behind the one controlling its motion (Fig. 3A)(*33*). When the first helicase reaches the linker at the end of the DNA section, it can no longer process and falls off. The DNA-peptide conjugate is then immediately pulled back into the nanopore such that the queued helicase, which is still bound to the DNA, takes control as the new anchoring enzyme. This effectively ‘rewinds’ the system and reinitiates a new independent read of the peptide sequence. The numbers of re-reads on the same single peptide can be very large: Fig. 3A shows an example of a raw data trace with 117 re-readings on a single peptide containing the G-substitution. We observed a typical rewinding distance of approximately 17 helicase steps, commensurate with a rewinding by a distance of ~17 amino acids or ~9 DNA bases, which indeed is the number of bases that is bound within the controlling helicase(*28*). Of the 117 re-reads in Fig. 3B, 45 re-reads stepped back far enough to provide a full re-read of the variant site.

**Figure 3:**
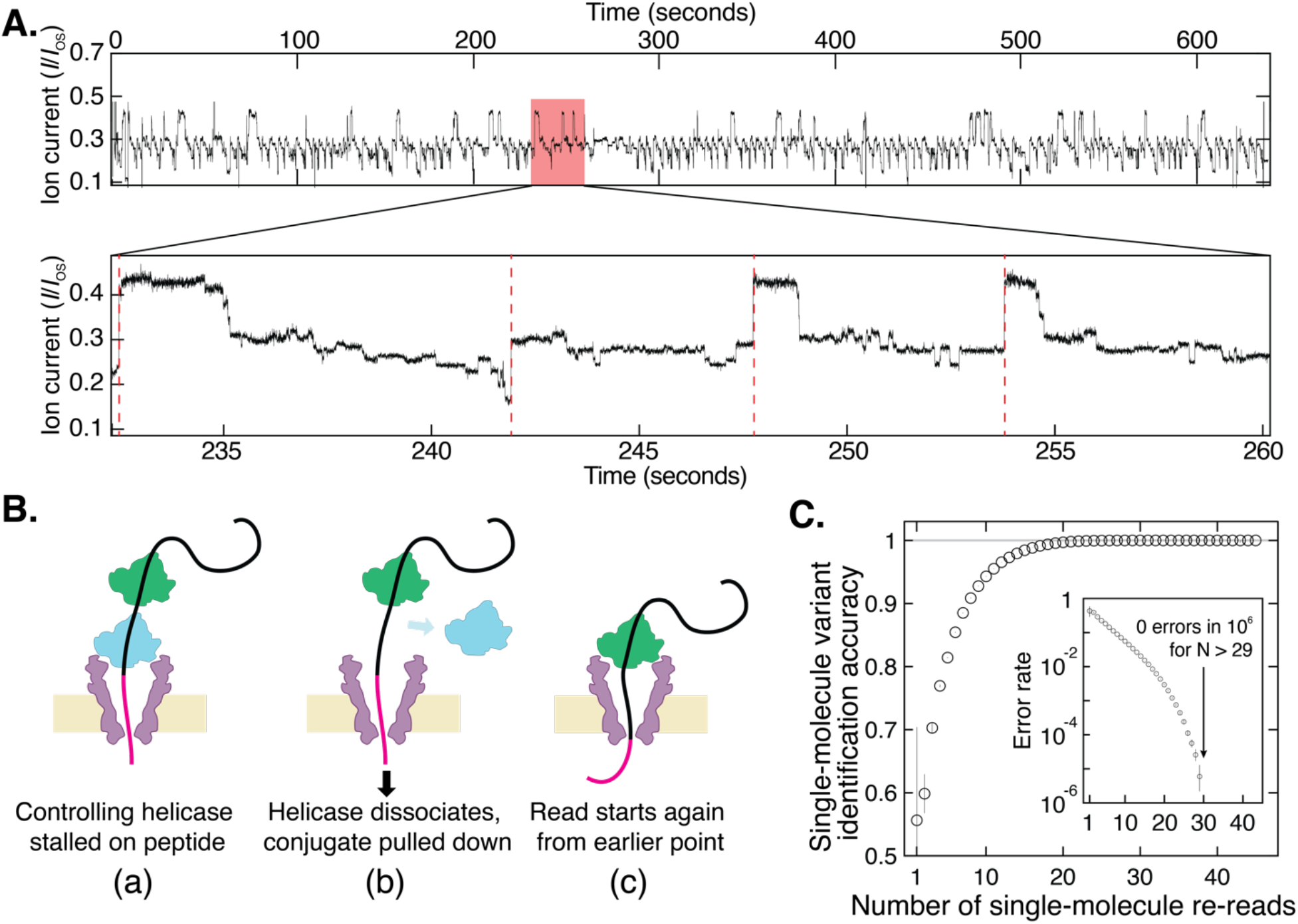
Re-reading of a single peptide sequence. (A) Highly repetitive ion current signal corresponding to numerous re-reads of the same section of an individual peptide (in this case, the G-substituted variant). The expanded plot below shows a region that contains four rewinding events (red dashed lines), where the trace jumps back to level 52 ± 2 of the consensus displayed in Fig. 2A. (B) Re-reading is facilitated by helicase queueing, where (a) a second helicase binds behind the primary helicase that controls the sequencing, re-reading starts when (b) the primary helicase dissociates, and (c) the secondary one becomes the primary helicase that drives a new round of sequencing. (C) By using information from multiple re-reads of the same peptide, the identification accuracy can be raised to very high levels of fidelity. These results indicate that with sufficient numbers of re-reads, random error can be eliminated and single-molecule error rate can be pushed lower than 1 in 106 even with poor single-pass accuracy. Inset is a logarithmic plot of the error rate = 1 − accuracy.

As Fig. 3C shows, we observe a marked improvement of the read accuracy with an increasing number of re-reads. To quantify the increase in the accuracy of the readings as a function of the number of re-readings, we generated randomly chosen subsets of the 45 measured re-reads and computed the identification accuracy with *N* re-reads as the fraction of subsets containing *N* re-reads that yielded the correct consensus identification (Supplementary Note 8). A striking improvement is observed upon re-reading: even when single reads are limited to as low as ~50% accuracy due to a partial coverage of the variant site, the re-reading method allows single molecules to be identified at extremely high levels of confidence, very close to 100%. As the inset to Fig.3C shows, the error rate very strongly decreases with the number of re-reads, yielding an undetectably low error rate (< 1 in 10^6^) when using more than ~30 re-reads of an individual peptide. Analysis on re-read traces from other variants yielded similar results (Supplementary Fig. S6).

The method described here provides proof of concept of a new approach for single-molecule protein sequencing, and is particularly powerful because of the re-reading mode of operation that eliminates primary nanopore sequencing errors. Transforming this into a technology capable of *de novo* protein sequencing will require taking reads of a broad library of peptides to build a map relating amino acid sequences to ion current levels, in order to assess the discriminatory power and accuracy of the technique across all 20 amino acids and many post-translational modifications. A range of heterogeneously charged peptides with neutral polar, nonpolar, negative, and positive amino acids can be explored (sample reads shown in Supplementary Fig. S8), for which, if needed, the MspA pore can be engineered to provide stronger electro-osmosis. The read length intrinsic to the technique, approximately 25-30 amino acids depending on the length of the DNA-peptide linker, does allow application of this method to many biologically relevant short peptides, such as 8-12 amino acid MHC-binding peptides(*34*). Additionally, this finite read length still represents a significant improvement over the small size of fragments used in mass spectrometry or Edman degradation, and protein fragmentation and shotgun sequencing methods similar to those used in traditional protein sequencing can naturally be applied to this new technique. Technical modifications such as using a variable-voltage control scheme (*29, 35*) have been shown to significantly improve the accuracy of DNA sequencing, and the physical principle of this is equally applicable to peptide sequencing (Supplementary Note 9).

Our protein re-reading method can be used immediately with any existing nanopore sequencing platform capable of accommodating MspA (e.g. the commercial MinION system), changing only the sample preparation and data analysis without requiring any re-engineering of the device. Furthermore, the method retains the features that enabled the success of nanopore DNA sequencing: low overhead cost, physical rather than chemical sensitivity to small changes in single molecules, and the flexibility to be re-engineered to target specific sequencing applications. Overall, our findings comprise a promising first step towards a low-cost method capable of single-cell proteomics at the ultimate limit of sensitivity to concentration, with a wide range of applications in both fundamental biology and the clinic.

## Acknowledgements

We thank Prof. Jens Gundlach and his lab members at University of Washington for providing the MspA nanopore and for sharing key pieces of software, and we thank Foteini Mentzou and Xin Shi for assistance with data collection, Eli van der Sluis for Hel308 purification, and Jaco van der Torre for his helpful advice on DNA construct preparation. This work was funded by NWO-I680 (SMPS) and supported by the NWO/OCW Gravitation programs NanoFront, the ERC Advanced Grant 883684, and the EC Marie Skłodowska-Curie action Individual Fellowship 897672. A.A. and J.L acknowledge support from NGHRI through grant R21-HG011741 and supercomputer time provided through XSEDE allocation MCA05S028 and the Blue Waters petascale supercomputer system (UIUC).

## Author Contributions

H.B. and C.D. conceived of the protein sequencing method. H.B. and A.K. conducted nanopore experiments and analyzed data. H.B. developed additional analysis code. J. L. and A. A. designed and conducted MD simulations. All authors discussed experimental findings and co-wrote the manuscript.

## Competing Interests

TU Delft has filed a patent application on technologies described herein, with H.B. and C.D. listed as inventors.

## Data and Materials Availability

All data and custom code used in this paper will be made available for download.

