## Supplementary Information for "Infinite re-reading of single proteins at single-amino-acid resolution using nanopore sequencing"

### Supplemental information for “Infinite re-reading of single proteins at single-amino-acid resolution using nanopore sequencing”

Henry Brinkerhoff, Albert S. W. Kang, Jingqian Li, Aleksei Aksimentiev, and Cees Dekker

July 13, 2021

### Contents

**1 Materials and Methods** **2**

**2 DNA-peptide hybrid construct design and assembly** **2**

**3 DNA ion current prediction** **4**

**4 Event selection** **4**

**5 Ion current level identification and filtering** **4**

**6 Consensus generation** **4**

**7 Variant identification** **8**

**8 Re-read analysis** **8**

**9 Variable voltage reads** **8**

**10 Heterogeneously charged peptide reads** **10**

**11 Molecular dynamics simulations** **10**

### List of Tables

S1 Sequences of DNA oligos and DNA-peptide hybrid oligos. . . . 3

### 1 Materials and Methods

Nanopore experiments were carried out as in previous work [6, 7, 20-24], on custom U-tube nanopore experimental devices. Experimental buffer consisted of 400 mM KCl, 10 mM $MgCl_2$, and 10 mM HEPES free acid at pH 8.00 ± 0.02. MspA was a kind gift from the laboratory of Jens Gundlach at the University of Washington, originally expressed by Genentech. Hel308 plasmid was obtained from Genscript (Cat. No. SC1849), and was expressed in-house using standard techniques. DNA-peptide conjugates and DNA oligos were obtained from Biomers. DPhPC lipid suspended in chloroform was obtained from Avanti. All experiments were performed at room temperature (21±1 ºC).

Nanopore ion current was recorded at 50 kHz sampling frequency with an Axopatch 200B patch clamp amplifier, and filtered with a 10 kHz 4-pole Bessel filter. Experiments were controlled through a National Instruments X series DAQ and operated with custom LabVIEW software. Data analysis was performed in Matlab. Preprocessing, data reduction and filtering, alignment and variant identification were performed using custom Matlab software described in the supplemental information and in previous work[6, 23].

Measured levels (red) in main text Figure 1D,E were identified by hand. Predicted levels (blue) were drawn from a 6-mer map of base sequence to DNA developed in previous work[23]. The highlighted linker and peptide sections were identified based on the length of the DBCO linker estimated from its chemical structure, as well as the consensus reads shown in figure 2A, where the variation resulting from the substitution determines the location of the substitution site.

The construction of the consensus reads in main text Figures 2A and B is described in Supplemental Information §6. Main text Figure 2C was generated by choosing the maximum likelihood variant based on a hidden Markov model alignment to each of the three variant consensuses, with the percentage calculated as (number of reads of variant X identified as variant Y)/(total number of reads of variant X), such that each row of the matrix sums to 100%. MD simulation methods used to generate main text Figures 2D-H are described in full in Supplemental Information §11.

To reliably obtain re-reads, helicase concentration was increased to $\approx 1$ $\mu$M. The segmentation and identification of re-reads used to generate the accuracy values in main text Figure 3C is described fully in Supplemental Information §8.

### 2 DNA-peptide hybrid construct design and assembly

The DNA-peptide hybrid constructs (main text Figure 1A) used to collect the bulk of the data were constructed of four components:

1. The template (variants “D22-”, “W22-”, “G22-”, and “hetero template” in Table S1), which consists of a 30-base nucleotide sequence attached at the 5’ end to the C-terminus of a 25-amino acid peptide by an azide-DBCO-C5 linker (Supplemental Figure S1A). This strand is pulled into the nanopore electrophoretically, and is read by the sequencer.
2. The complement (“complement gen2” in Table S1), which is complementary to part of the template strand and serves three functions: (a) a 3’ cholesterol allows it to associate with the bilayer, increasing the frequency of DNA-pore interactions; (b) a 5’ overhang provides a sticky end to attach the template extender; and (c) on the hybridized construct, it blocks the Hel308 enzyme (which has poor helicase processivity) from processing along the template and using ATP, until the template enters the pore and the complement is sheared off by MspA.
3. The 50-base template extender (“template extender gen2”), which binds to the sticky 5’ end of the complement, and extends with its own 10-base 3’ sticky end which acts as a binding site for Hel308. The ligation of this extender is necessary to increase the length of the DNA-peptide hybrid, which is only commercially available in lengths too short to be efficiently captured by the pore.
4. The staple (“staple”), a 10-base oligo complementary to both the template and the extender, which enables the efficient ligation of the two oligos. The staple is used to prepare the construct, but once assembled has no functionality in the construct, and like the complement is sheared off upon capture by MspA.

The exact sequences used are provided in Supplemental Table S1. To assemble the constructs, equal amounts of template, staple, and template extender were mixed and annealed in a thermocycler by heating to 95 ºC for 2 min and letting them cool down slowly to room temperature. The mixture was then incubated for 18h at 16 ºC with 400U of T4 DNA ligase (NEB, M0202T) in 1X of the manufacturer-provided buffer solution. Next, a 1.1X excess of the ligated construct was mixed with the complement at $> 1$ $\mu$M concentration of each and annealed.

Some reads (those labeled “biomers1cholext” as opposed to “biomers1cholext2” or “biomers1cholext2nohp” in the Supplementary Data) used an earlier version of the construct, in which the complementary strand itself (“complement gen1” in

| Oligo name | Sequence |
|---|---|
| D22 template | [N-term] DEDEDEDEDEEDDEDEDDEEEDDDD [C-term] (linker) 5’ TTACTGAAGTCTCACGTGCCTGGTATATTA 3’ |
| W22 template | [N-term] DEDEDEDEDEEDDEDEDDEEEWDDD [C-term] (linker) 5’ TTACTGAAGTCTCACGTGCCTGGTATATTA 3’ |
| G22 template | [N-term] DEDEDEDEDEEDDEDEDDEEEGDDD [C-term] (linker) 5’ TTACTGAAGTCTCACGTGCCTGGTATATTA 3’ |
| hetero template | [N-term] DDDDDDDDDDDDDDYAVEGRDLTLS [C-term] (linker) 5’ TTACTGAAGTCTCACGTGCCTGGTATATTA 3’ |
| complement gen1 | 5’ TGATCAATTCACTGTGGATGTAATATACTTTTTTTTTTTTTTTTTTTTTTTTTTTTTTTT 3’ (cholesterol) |
| complement gen2 | 5’ GATGTAGAATTTTTTTTTTTTTTTTTTTTT 3’ (cholesterol) |
| template extender gen1 | (phosphate) 5’ CATCCACAGTGAATTGATCAGGTCGTAGCC 3’ |
| template extender gen2 | (phosphate) 5’ CATCCACAGTGAATTGATCATTATGACGTTATTCTACATCGGTCGTAGCC 3’ |
| staple | 5’ GGATGTAATAGC 3’ |

Table S1: Sequences of DNA oligos and DNA-peptide hybrid oligos.

Table S1) was used to assemble and ligate the template to an extender (“template extender gen1”). This construct is illustrated in Figure S1B. This resulted in both a shorter free length of ssDNA/peptide lowering the rate of template capture by MspA, and the longer cholesterol tether being frequently captured and occupying the pore while excluding the template. These negative impacts on throughput led us to the revised construct described above. The end of the DNA sequence and the peptide sequence read in this earlier generation was identical to that in the later reads, so the signal from the region of interest discussed in the paper was unchanged.

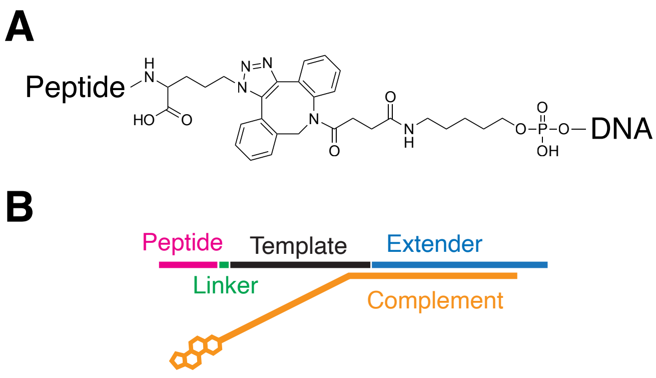

Figure S1: Details of sequencing constructs. (A) The chemical structure of the DBCO-Azide linker used in the DNA-peptide hybrids.(B) First-generation sequencing construct that was used in early data acquisition, which ultimately was replaced by the one shown in main text Figure 1A.

### 3 DNA ion current prediction

Following previous work *(29)*, ion currents for the DNA section in main text Figure 1E were predicted using an empirically derived 6-mer map, converting each 6-base subsequence into ”pre-” and ”post-” ion current states corresponding to the two substeps of Hel308 per DNA base.

The construction of the 6-mer map from measurements of genomic DNA is described in Noakes 2019, supplemental material §7. Briefly, ion current measurements corresponding to each 6-mer pre- and post-state were obtained from kilobase or longer reads of genomic $\lambda$ phage and $\Phi$ X174 viral DNA. The ion currents in the map are the mean of the set of ion currents assigned to each state, and the uncertainty in the ion current is the standard deviation in that set of ion currents.

### 4 Event selection

Candidate events were identified with a simple thresholding algorithm and filtered by duration, keeping only those blockages longer than 1 second. The candidate reads were then inspected by eye, and only reads matching the DNA prediction in their first part, and containing further enzyme stepping behavior after the end of the DNA were retained. Reads containing significant amounts of MspA gating or spurious noise, or reads with fewer than 12 observed levels were also rejected. These criteria are illustrated in Figure S2. As visible in figure S2 as well as figure S3, the ion current sequence of reads is highly reproducible, but subject to the usual random error intrinsic to nanopore reads, much of which may be systematically removed. Because some analyses rely on separate analysis of the peptide section, we identified the peptide section as beginning at consensus level 49.

### 5 Ion current level identification and filtering

To segment the data for consensus refinement through expectation maximization and for blinded variant identification, we used a change point detection algorithm exactly as described in previous work (Noakes 2019 Supplemental Information §11 *(29)*; originally described in Wiggins 2015 *(36)*). A sample trace with automated level finding indicated is shown in Figure S4.

The further filtering steps described in Noakes 2019 were also applied to the resultant level sequences. First, a state filter was applied to excise levels that were too short ($<$ 2 ms) or significantly outside the bounds of the consensus currents ($I/I_{\mathrm{OS}} < 0.25$ or $I/I_{\mathrm{OS}} > 0.5$). These filters serve to eliminate a significant number of spurious states resulting from noise spikes or mid-event MspA gating. Next, a backstep recombination filter was applied using the algorithm of Noakes 2019, Supplementary Information §5.2, in order to eliminate the bulk of helicase backsteps. The recombination filter, which relies on comparing levels in an event to other nearby levels in the same event, is more accurate than accounting for a large number of backsteps in the alignment algorithm, because read-to-read error, which can impact the scoring of an alignment to reference, does not affect the matching of observed states in a self-comparison.

### 6 Consensus generation

Consensus reads were generated through a customized Baum-Welch algorithm, a type of expectation maximization (EM) for the hidden Markov model. The EM algorithm, described fully in previous work *(29)* (Noakes 2019, Supplement §7.4) is as follows:

1. Solve the hidden Markov model using a maximum-*a posteriori* likelihood (MAP) algorithm to assign likelihoods that each of the ion current levels in each read were produced by a particular true template position within the constriction (helicase step number).

2. If the change in log likelihood of the HMM solution is greater in magnitude than a threshold (in our case $10^{-3}$), continue. Otherwise, reject the latest consensus sequence and terminate.

3. Compute a new mean and uncertainty in ion current value for each HMM state using an average of the measured values weighted by the probability that each value was assigned to that state.

The EM algorithm requires an initial guess at an HMM in order to begin. To seed the EM algorithm with an initial set of HMM observation probabilities, a selection of typical reads of each construct were cross-compared by eye to identify the unique ion current states and put them in the correct order, while eliminating single-read errors like spurious states, missed states, or enzyme backsteps. The result was a set of aligned sequences of levels, where each level is characterized by a mean, standard deviation and number of measurements included.

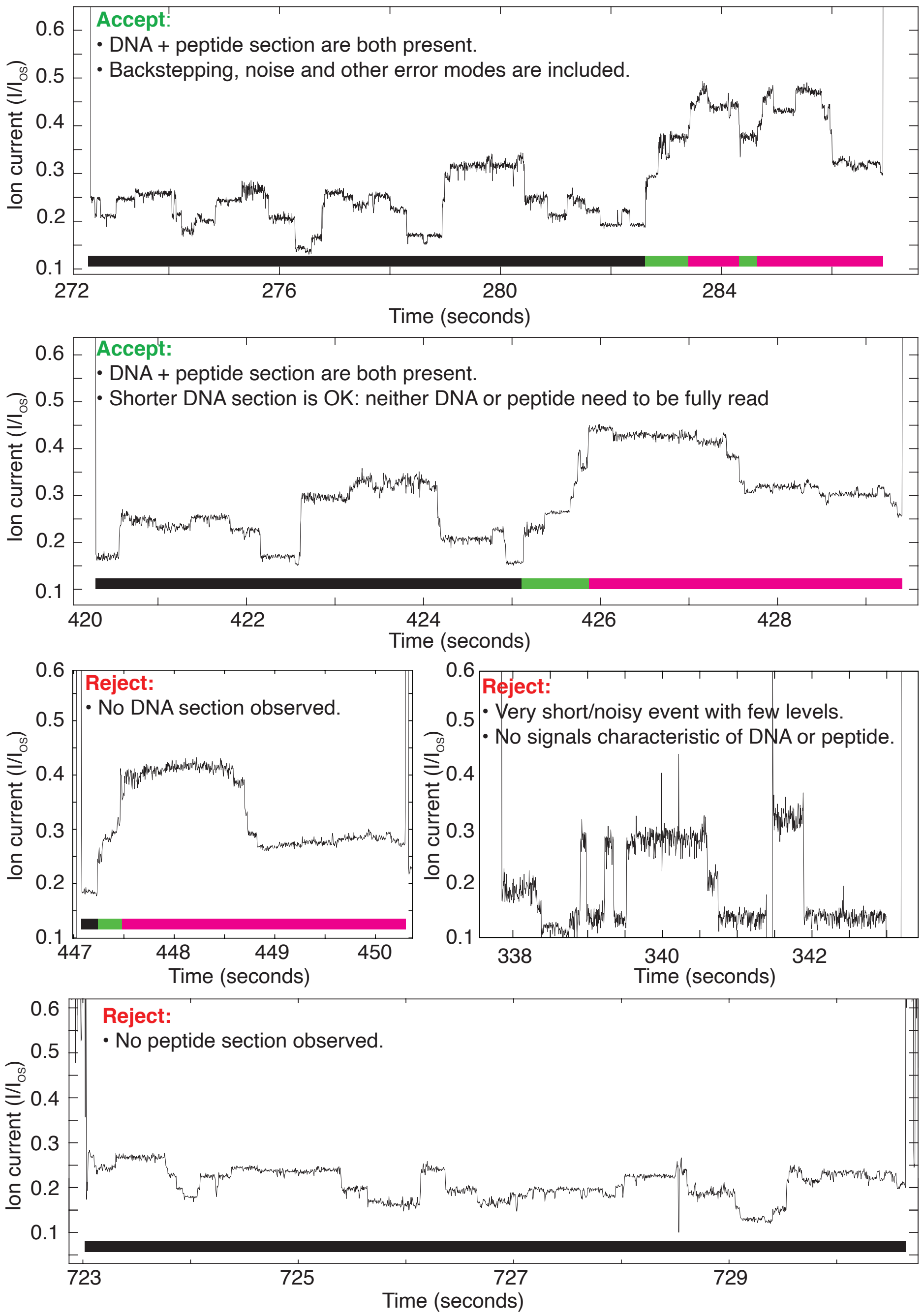

Figure S2: Examples of accepted and rejected events. Colored bar at the bottom indicates which portion of the conjugate polymer is in the constriction of MspA: black = DNA, green = linker, magenta = peptide.

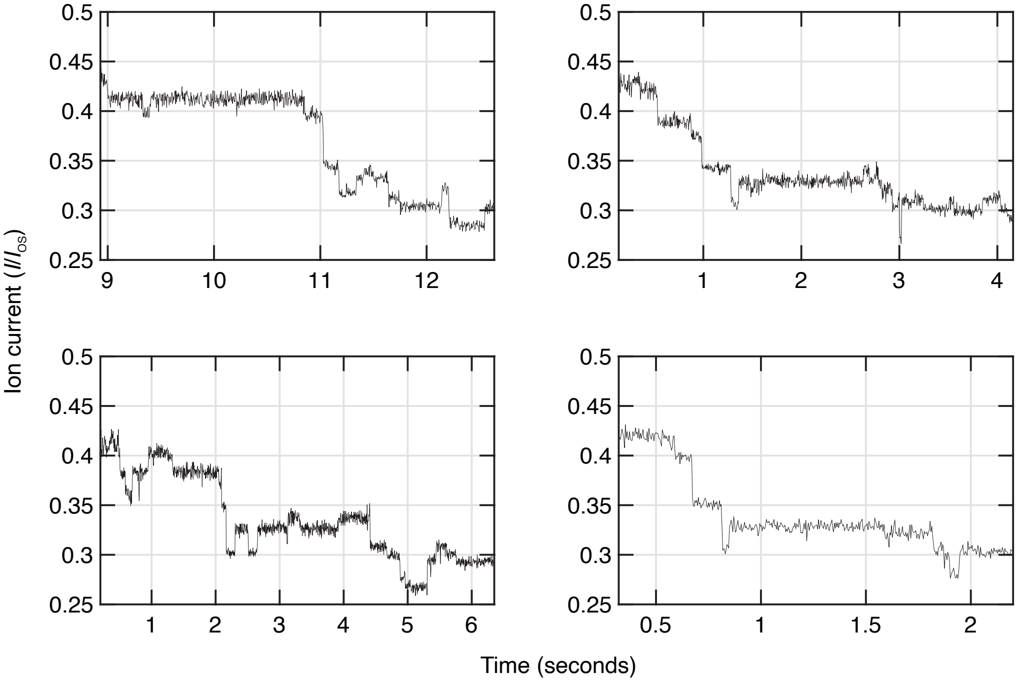

Figure S3: Reproducibility of ion current sequences. A random selection of the peptide section of W-variant reads used in the paper, demonstrating the consistency of the observed ion current levels. Different reads contain apparent variation: they differ considerably in terms of the durations of individual helicase steps, noise sometimes makes it difficult to segment steps, very short states are sometimes apparently missing from reads, and numerous helicase backsteps are apparent. However, the underlying sequence of ion current means is highly reproducible, and the random errors may be accounted for in the process of backstep removal (Supplementary §5) and read alignment to consensus (Supplementary §7), or eliminated through re-read analysis (main text Figure 3).

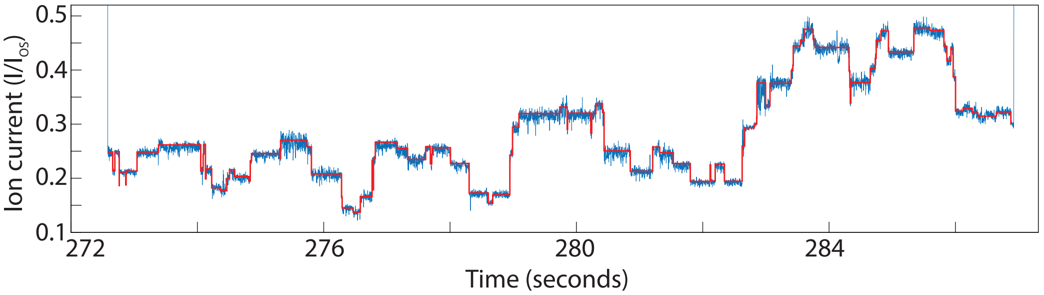

Figure S4: Typical performance of information-based level finder on nanopore data.

Different nanopore reads may vary by an overall scale in ion current due to variations in buffer salt concentration caused by evaporation and due to day-to-day variations in temperature *(7)*. Therefore, reads must always be calibrated by applying an appropriate scale $m$ to all ion currents they contain. To find the maximum likelihood estimators for $m$ for all $N$ aligned sets of reads of length $L$, we want to minimize the total error between reads

$$\hat{\boldsymbol{m}} = \arg\min_{\boldsymbol{m}} \sum_{k=1}^{L} \sum_{i,j=1}^{N} \begin{cases} \frac{(m_i x_{ik} - m_j x_{jk})^2}{\delta x_{ik}^2 + \delta x_{jk}^2} & \text{if } x_{ik} \text{ and } x_{jk} \text{ both exist,} \\ 0 & \text{otherwise.} \end{cases} , \tag{1}$$

where $x_{ik}$ and $\delta x_{ik}$ are the mean and uncertainty in the mean of state $k$ in read $i$, and we have made the approximation that the uncertainties do not scale with the ion currents when the near-unity calibration is applied. However, this optimization still leaves one degree of freedom: the sum is invariant if every read is subject to the same overall scale. To eliminate these ambiguities, we choose by convention to also include the requirement that $\frac{1}{N}\sum_i m_i = 1$. We end up with a full rank linear system of equations, which can be easily solved for $\hat{\boldsymbol{m}}$. The scales are applied to the reads, and a consensus mean, standard deviation, and uncertainty are then computed as

$$\bar{x}_k = \frac{1}{N}\sum_{i=1}^{N} x_{ik}, \tag{2}$$

$$\bar{\sigma}_k = \frac{1}{N}\sum_{i=1}^{N} \sigma_{ik}, \tag{3}$$

$$\delta\bar{x}_k = \sqrt{\frac{1}{N}\sum_{i=1}^{N} \delta x_{ik}^2 + \frac{1}{N}\sum_{i=1}^{N} (x_{ik} - \bar{x}_k)^2} \tag{4}$$

where $x_{ik}$ now refers to an element of a calibrated read.

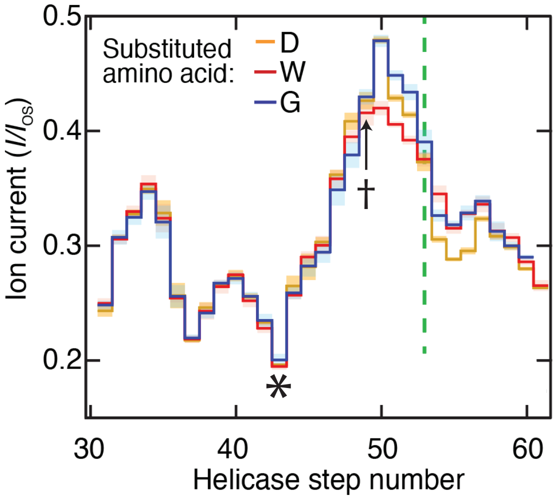

Figure S5: Important consensus levels. Level 43, marked with an asterisk *, was used to calibrate all reads with an overall scaling of ion current. This level was identified in raw data as the locally minimal ion current level preceding the maximum ion current level in the trace. Level 49, marked with a dagger †, marked the beginning of the peptide section used for analysis. This level was identified in raw data as the level immediately preceding the maximum ion current level in the trace.

Next, to ensure cross-construct calibration consistency, the same procedure is carried out to find an optimal scale and offset for each of the three handmade consensuses using only the DNA section of the reads. Since the DNA section is known to be identical across different reads, we replace the DNA section in each consensus with its mean across all three variants. The means and standard deviations of the three calibrated consensuses are used as initial guesses for the EM algorithm.

Calibration scales also need to be found for every read, including those not used in the handmade consensus generation. These reads were calibrated straightforwardly by choosing a scale such that the ion current of level 43 (see Figure **??**, asterisk *) matched the value of that state in the average of the three consensuses. This level was chosen because it was easily identifiable, relatively low in noise, and was present in every analyzed measurement with both DNA and peptide sections due to its position between the two regions.

With a set of properly calibrated reads and an initial guess for the consensus, we updated the peptide section of each consensus by running the EM algorithm to convergence using a randomly chosen subset of the peptide section of 20 of each variant's reads, and thus arrived at the three consensuses used to classify the reads to produce Figures 2C and 3C in the main text.

### 7 Variant identification

When identifying variants, we used those calibrated reads randomly reserved from inclusion in the consensus generation. Using a Viterbi algorithm accommodating both forwards and backwards steps, unobserved steps, spurious ion current levels, and over-segmented levels (described in previous work: Laszlo 2014*(7)*, supplementary note 2), the peptide section consisting of levels 49 (see Figure **??**, dagger †) to either termination or a rewinding event, of each read was aligned and assigned likelihood scores, and the highest scoring read was determined to be the variant for that read.

### 8 Re-read analysis

Re-reads were reliably obtained by increasing helicase concentration to an excess of 1 $\mu$M. At lower helicase concentrations, events with 2-4 re-reads were seen occasionally.

For the re-read analysis in main text Figure 3, a particularly long read of the G22 template containing approximately 117 rewinding events was parsed by hand to separate each separate re-read (Figure S6A). They were then processed using the level segmentation, filtering and backstep removal described in Supplementary §5. Only re-reads with at least one ion current level greater than 0.35 $I/I_{\text{OS}}$ were included in the analysis shown in main text Figure 3C. This restriction was applied in order to ensure that the read re-wound at least to consensus level 53, covering at least half of the ion current level sequence affected by the substitution, and omit attempts to identify the variant from reads of a section not containing the substitution. 45 re-reads were ultimately included in the final analysis. Each re-read was assigned a likelihood of being drawn from each of the three variants, as described in supplemental §7, using a consensus trained on all 216 single-read events (available in the Supplementary Data).

The accuracies in main text Figure 3C were computed as follows: For each integer value of $N = 1$ to 45, $10^6$ randomly selected subsets of $N$ reads were generated. For $N = 1$, 2, 43, 44, and 45, the number of possible subsets is smaller than $10^6$, so the random sampling closely reproduces the results of analyzing all possible subsets of these sizes.

For each subset, the variant likelihoods of each re-read in the subset were multiplied and then normalized to sum to 1. The maximum-likelihood variant was then chosen. The $N$-re-read accuracy was defined as the proportion of the $10^6$ subsets of size $N$ whose maximum-combined-likelihood variant was the true (G) variant for the read. For example, to estimate the accuracy from 10 re-reads, we randomly selected 10 of the re-reads in the event. We calculate the likelihood that each of the 10 reads was of each variant, and multiply together the 10 values for each variant. The variant with the largest product is the identification. This is repeated $10^6$ times, and the number of 'G' identifications was divided by $10^6$ to obtain the fractional accuracy. Single-pass accuracy in the re-read data (the N=1 data point in main text Figure 3C) is lower than that reported in main text Figure 2C because re-reads often only partially cover the variant site, resulting in some loss of sequencing information.

This approach was carried out with further reads, including many shorter than the one shown in the main text, and the results are summarized in figure S6B.

In future work, we expect that even better results can be obtained by first generating a consensus sequence of ion currents, and using that signal to classify variants or sequence, rather than simply combining match likelihoods, because this in principle allows for convergence on an easily recognizable consensus sequence much faster, typically with fewer than 10 re-reads.

### 9 Variable voltage reads

In previous work*(29)*, we showed that by driving a nanopore experiment with a time-varying voltage, and thereby measuring a conductance-voltage curve at each enzyme step instead of only a mean ion current, we obtained significantly more detailed information about the DNA being sequenced and single-read sequencing accuracy increased dramatically. The principle of this method is applicable to any polymer analyte whose mean position in the pore depends on the pulling force applied by

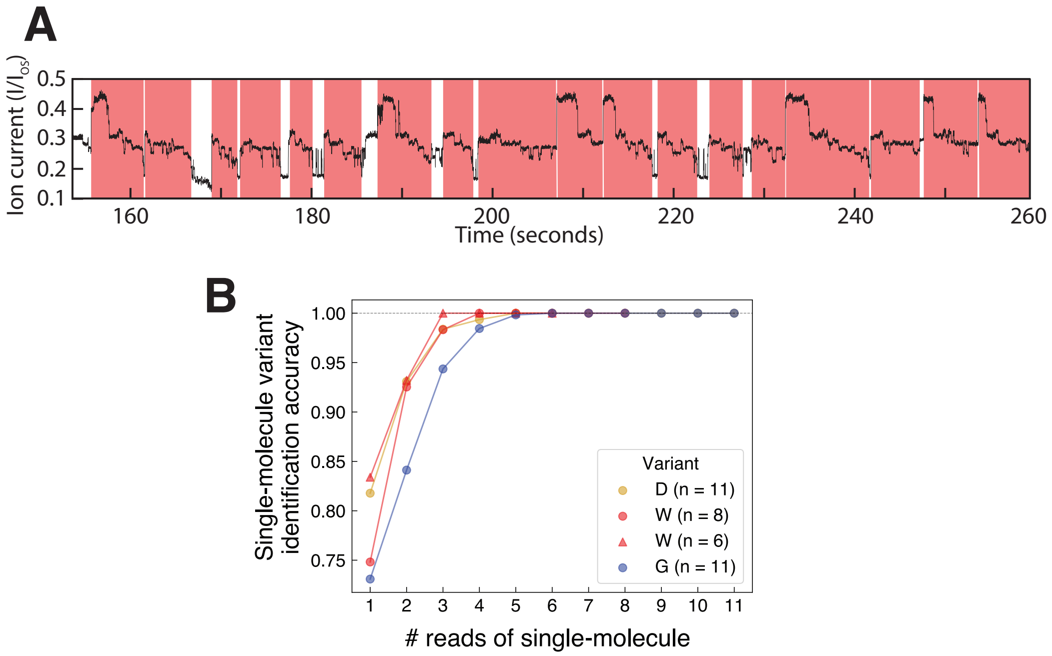

Figure S6: Re-read analysis. (A) Re-read segmentation. Each identified independent re-read is bounded in a red highlighted region. Re-read ends were marked at approximately consensus level 60, or at the end of the re-read if rewinding occurred before level 60. Re-read beginnings were marked when the current returned to a previously visited level more than 3 steps back, in order to avoid representing normal helicase backsteps as very short re-reads. (B) Plots similar to main text Figure 3C for several events with various numbers $n$ of total re-reads. Color indicates true variant: gold = D-variant, blue = G-variant, red = W-variant. Two multi-read events, one G-variant with n = 4 rereads, and one D-variant with n = 9, yielded 100% accuracy for all re-reads in the event and are omitted from the plot for clarity.

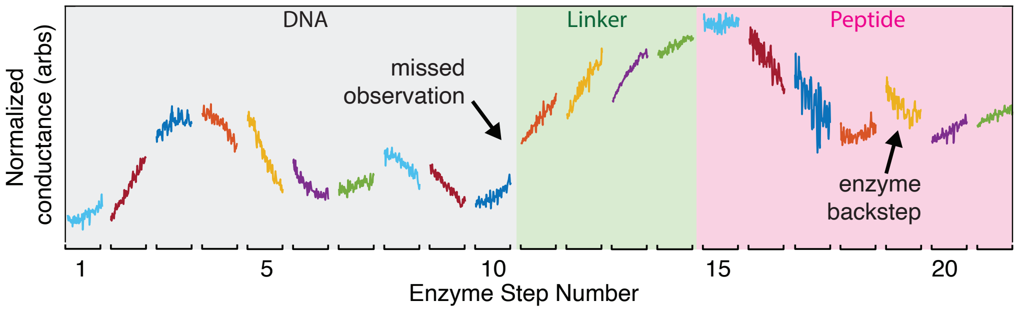

Figure S7: Sample variable-voltage read. This variable-voltage read covers both the DNA and D-variant peptide parts of the sequence. Compare the trends in conductance to the trends in ion current seen in main text Figures 1D, 1E, and 2A. Major qualitative features yielding improved sequencing fidelity in DNA reads are retained in reads of the peptide, including the smooth character of the conductance-voltage curve at each enzyme step, and the rough continuity of a forward-stepping read allowing us to infer where backsteps or missed observations occur in a read. Colors are only to aid the eye in discerning separate ion conductance curves.

the voltage, including electrophoretically or electro-osmotically trapped peptides.

To assess the feasibility of future experiments using high-fidelity variable voltage sequencing, we obtained and processed variable voltage reads exactly following the methods of Noakes 2019. These reads display similar qualitative characteristics to variable voltage experiments with DNA: namely, the ability of each conductance-voltage curve to be fit well with a second-order polynomial, and rough continuity of the curves enabling backsteps and unobserved enzyme steps to be more readily identified (Figure S7). This suggests that in future work, using the variable-voltage sequencing method will allow for comparable improvements to peptide sequencing fidelity. In the current paper, we focused instead on the constant-voltage current stepping signals for clarity of communication.

### 10 Heterogeneously charged peptide reads

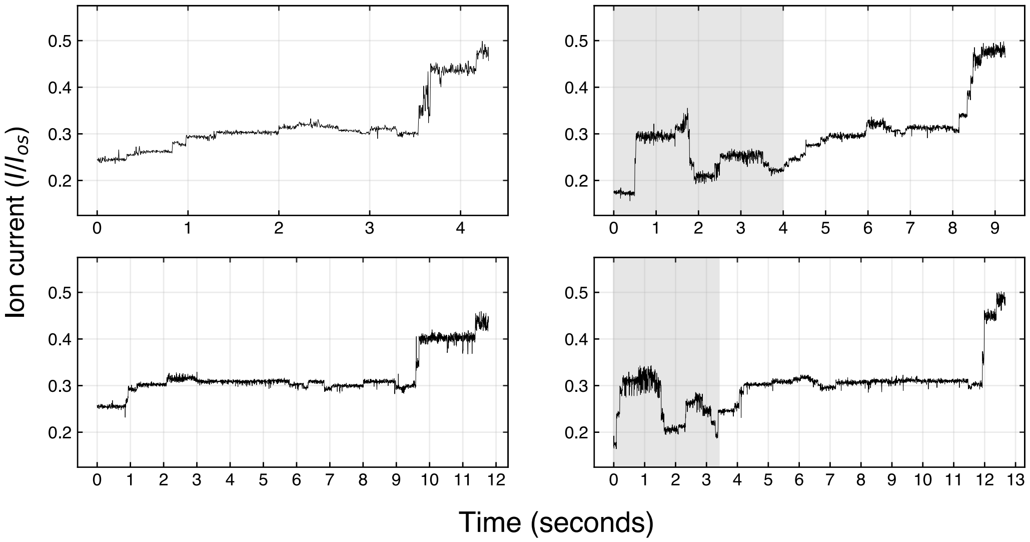

Figure S8: A selection of reads of heterogeneously charged peptides. Clear stepping can be seen, with a similar number of steps as observed in the negatively charged peptide experiments. The gray shaded regions in the right column reads correspond to the DNA portion of the read; compare to main text Figure 1E levels 1 through 44.

Our paper presents the principle of our approach using model peptides that are homogeneously charged. Preliminary data indicate that the method will also work with more heterogeneous template constructs ("hetero template" in Supplementary Table S1), which consist of a mixture of positive and negative charges as well as polar and non-polar neutral side chains. Indeed, first reads yielded clear and reproducible stepping ion currents (Figure S8). While further experimentation and analysis will be required to fully understand the ion current signals observed, these reproducible traces are evidence for the sufficiency of the electro-osmotic force to trap peptides in MspA for cases where a strong electrophoretic force is absent.

### 11 Molecular dynamics simulations

**General MD Methods.** All simulations were performed using the classical MD package NAMD *(37)*, periodic boundary conditions, and a 2 fs integration time step. The CHARMM36 force field *(38)* was used to describe proteins, dioctadeca-trienoylphosphatidylcholine (DPhPC) phospholipids*(39)*, TIP3P *(40)* water, and ions *(41)* along with the CUFIX corrections applied to improve description of charge-charge interactions *(42)*. RATTLE *(43)* and SETTLE *(44)* algorithms were applied to covalent bonds that involved hydrogen atoms in protein and water molecules, respectively. The particle mesh Ewald (PME) *(45)* algorithm was adopted to evaluate the long-range electrostatic interaction over a 1 Å-spaced grid. Van der Waals interactions were evaluated using a smooth 10–12 Å cutoff. Langevin dynamics were used to maintain the temperature at 295 K. Multiple time stepping was used to calculate local interactions every time step and full electrostatics every two time steps. The Nose-Hoover Langevin piston pressure control *(46)* was used to maintain the pressure of the system at 1 atm by adjusting the system's dimension. Langevin thermostat *(47)* was applied to all the heavy atoms of the lipids with a damping coefficient of 1 $ps^{-1}$ to maintain the system temperature at 295 K.

**MD Simulations of MspA Nanopores Containing Peptides.** An all-atom model of reduced-length MspA was constructed as described previously *(30)* to include residues 75–120 of the full-length protein, merged with an 8 × 8 $nm^2$

patch of DPhPC bilayer and solvated with 0.4 M KCl electrolyte, a system of approximately 39,500 atoms. Thirty-two aspartate residues were replaced by asparagines or arginine to create the D90N/D91N/D93N/D118R mutant used in experiment. Nine additional systems were constructed to have a 23-amino acid polypeptide strand placed inside the nanopore to span through the nanopore constriction, differing by the amino acid sequence and the location of the single amino acid substitution relative to the constrictions, see Figures S9–S11 for details. The peptides were built to have a stretched conformation characterized by the end-to-end distance of approximately 65Å.

Following assembly, each peptide system was minimized in 2,000 steps using the conjugate gradient method and then equilibrated for 45 ns at a constant number of atoms, pressure, and temperature (NPT) ensemble performed while keeping the ratio of the systemÕs size along the place of the bilayer constant. During the equilibration and in all subsequent simulations, a harmonic restrain ($k_{\mathrm{SPRING}} = 10$ kcal mol$^{-1}$ Å$^{-2}$) was applied to the C$_\alpha$ atom of top (C-terminal) residue of the peptide. Additionally, each C$_\alpha$ atom of the MspA protein was harmonically restrained ($k_{\mathrm{SPRING}} = 1$ kcal mol$^{-1}$ Å$^{-2}$) to its X-ray coordinate *(27)*. The systems were then simulated for 50 ns in a constant number of particles, volume and temperature (NVT) ensemble under a constant electric field $E = -V/L_z$ applied along the $z$-axis (normal to the membrane) to produce a transmembrane bias $V$; where $L_z$ is the dimension of the simulated system in the direction of the applied electric field *(48,49)*. For the NVT simulations, the systemÕs dimensions were set to the average dimensions observed within the last 5 ns of the restrained NPT equilibration.

To obtain a representative ensemble of peptide conformations within the nanopore, each of the seven peptide systems were simulated under a transmembrane bias of either 200 mV (G and D systems) or 600 mV (W systems) while moving the top residues of the peptide strand by 6 Å away from the constriction and back, four times over the course of 400 ns, Figures S9–S11. Forty five instantaneous configurations were chosen from the three simulations of the G and W systems producing an ensemble of conformations differing by the location of the amino acid substitution relative to the MspA constriction; five configurations were chosen from the single simulations of the D system. Each system was then simulated for 200 ns under 200 mV having the top residue of each peptide stationary restrained to its coordinate in the chosen instantaneous configuration. During these 200 ns simulations, the amino acid substitutions maintained their z coordinate within $-5$ to $+15$Å from the nanopore constriction, Figure S12. The blockade current analysis was done on these 50 trajectories.

The open pore system was minimized (2,000 steps) and equilibrated (45 ns in NPT) similar to the peptide systems. The open pore current was obtained from a 450 ns NVT simulation under a 200 mV bias.

**Ion Current Calculation.** Instantaneous ionic current was calculated as *(48)*

$$I(t) = \frac{1}{\Delta t l_z} \sum_{j=1}^{N} q_j (z_j(t + \Delta t) - z_i(t)), \tag{5}$$

where $z_j(t + \Delta t) - z_i(t)$ is the displacement of ion j along the $z$ direction during the time interval $\Delta t = 20$ ps and $q_{\mathrm{j}}$ is the charge of ion j. To minimize the effect of thermal noise, the current was calculated within an $l_z = 20$ Å thickness slab centered at the nanopore constriction (the slab spanned the entire simulation system in the $x$-$y$ plane). The instantaneous values of the ionic current were recorded simultaneously with the center of mass $z$ coordinate of the backbone of the amino acid substitution. For each amino acid substitution, the data from all trajectories were sorted according to the $z$ coordinate of the substitution in ascending order. The average value of the current and its standard error were computed using 2 Å bins along the $z$ coordinate.

**Calculations of Nanopore Volume.** To calculate the fraction of the nanopore volume available to conduct ionic current, we first computed the average number of bulk-like water molecules confined within the nanopore constriction for the open pore simulation. Bulk-like water molecules were defined as those located more than 2.5 Å away from any protein atoms. Previously, we found the number of bulk-like water molecules in the MspA constriction to determine the ionic current through MspA blockade by a DNA strand *(30)*. Following that, we calculated the instantaneous number of bulk-like water molecules in the nanopore constriction for every second frame of the same 50 MD trajectories that we used for the ionic current blockade analysis (23 for G, 22 for W and 5 for D systems, Figure S12). The nanopore constriction was defined as the inner volume of the nanopore located within 4Å along the $z$ axis from the center of mass of residues 90 and 91. The fraction of nanopore constriction volume occupied to conduct ionic current was obtained by dividing the number of bulk-like water molecules in the nanopore constriction blocked by the peptide by the number of bulk-like water molecules in the open pore constriction. For each amino acid substitution, the data from all trajectories were sorted according to the z coordinate of the substitution in ascending order. The average value of the volume fraction and the standard error were computed using 1 Å bins along the $z$ coordinate.

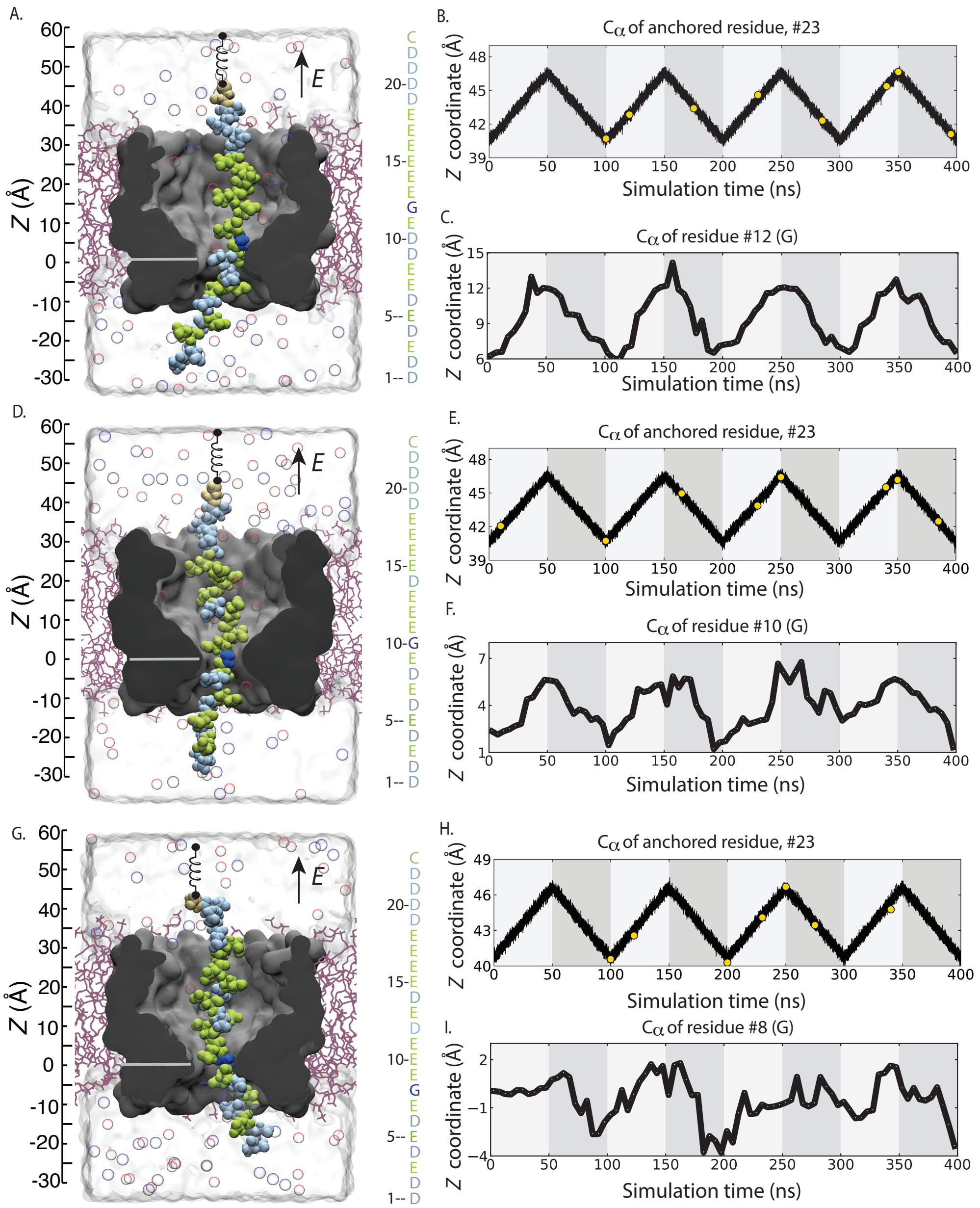

Figure S9: Preparation of initial configurations for the production simulations of the G system. (A–C) Starting from the pre-equilibrated configuration shown in panel A, the system was simulated using the all-atom MD method under a 200 mV bias while the $C_\alpha$ atom of the top residue (#23) was moved up and down by approximately 6 Å four times in 400 ns, panel B. Panel C shows the corresponding displacements of the G residue at position #12 by plotting to coordinate of the backbone $C_\alpha$ atom. The yellow circles in the trace in panel B show the initial configurations chosen for the production simulations, Figure S12B. (D–F and G–I) Same, but for different placement of the W mutation (at residue #10 and #8) relative to the pore constriction.

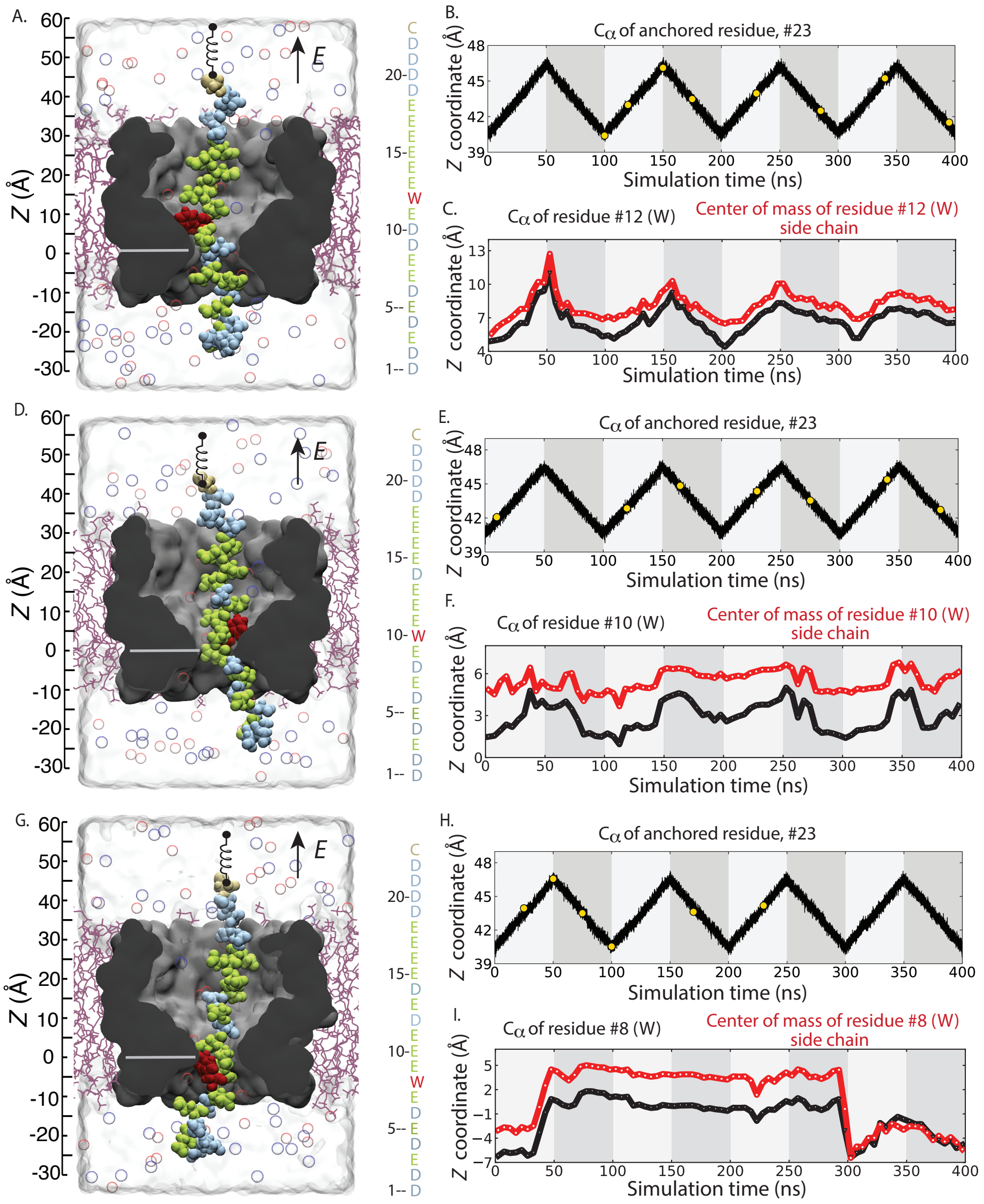

Figure S10: Preparation of initial configurations for the production simulations of the W system. (A–C) Starting from the pre-equilibrated configuration shown in panel A, the system was simulated using the all-atom MD method under a 600 mV bias while the $C_\alpha$ atom of the top residue (#23) was moved up and down by approximately 6 Å four times in 400 ns, panel B. Panel C shows the corresponding displacements of the W residues at position #12 separately for the backbone $C_\alpha$ atom and for the side chain center of mass. The yellow circles in the trace in panel B show the initial configurations chosen for the production simulations, Figure S12B. (D–F and G–I) Same, but for different placement of the W mutation (at residue #10 and #8) relative to the pore constriction.

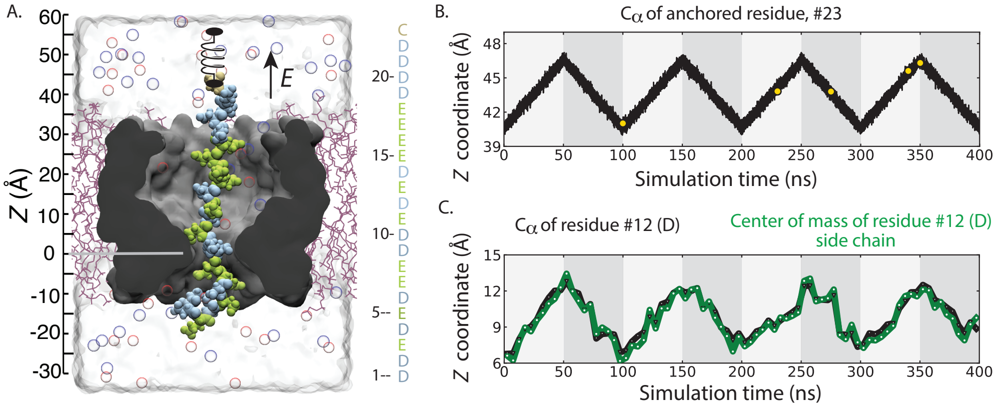

Figure S11: Preparation of initial configurations for the production simulations of the D system. Starting from the pre-equilibrated configuration shown in panel A, the system was simulated using the all-atom MD method under a 200 mV bias while the $C_\alpha$ atom of the top residue (#23) was moved up and down by approximately 6 Å four times in 400 ns, panel B. Panel C shows the corresponding displacements of the D residues at position #12 separately for the backbone $C_\alpha$ atom and for the side chain center of mass. The yellow circles in the trace in panel B show the initial configurations chosen for the production simulations, Figure S12C.

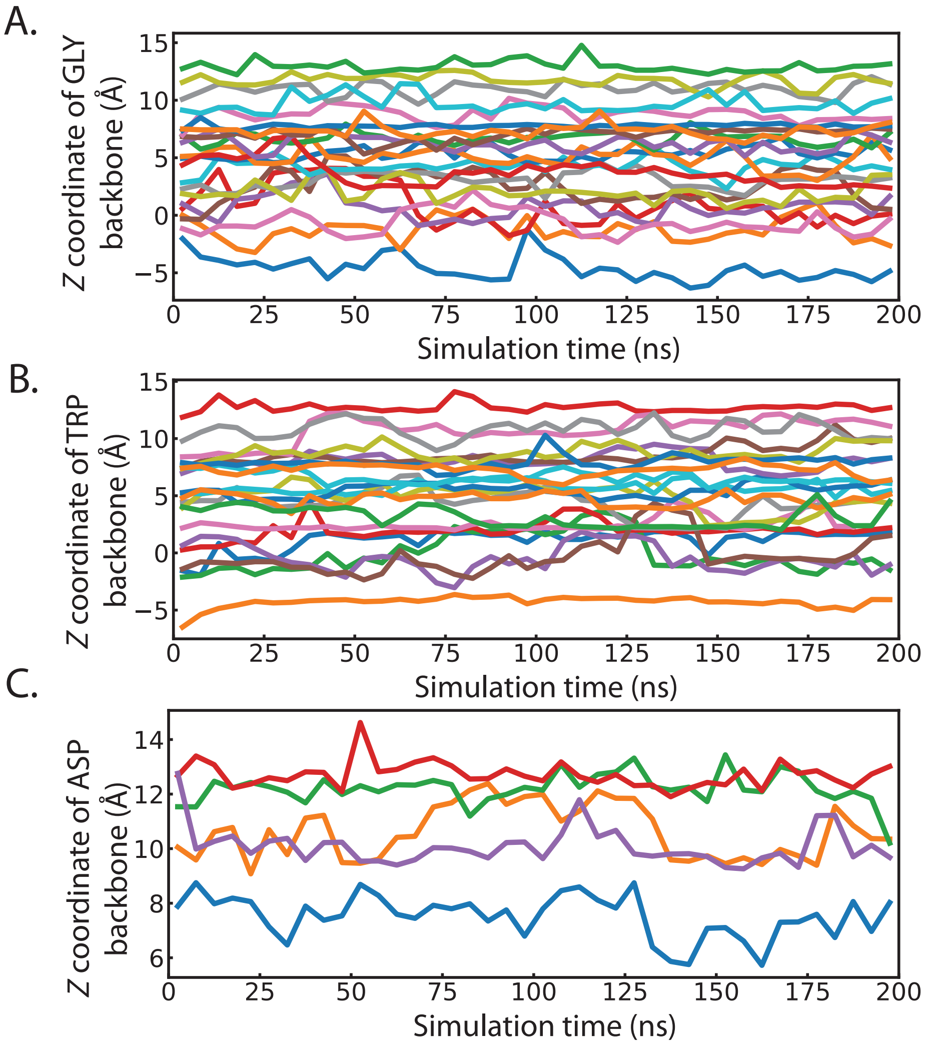

Figure S12: Production simulations of MspA–peptide systems. (A–C) Center-of-mass $z$ coordinate of a single amino acid backbone versus simulation time for 49 independent MD simulations carried out under a 200 mV bias. Data in panels A, B and C correspond to 23 G, 22 W and 5 D systems. The initial states for the MD simulations were chosen from the periodic displacement simulations featured in Figures S9–S11.
